# Downregulation of Gankyrin/PSMD10 affects cancer cell growth and proliferation in glioblastoma: *in vitro* and *in vivo* study

**DOI:** 10.1101/2024.12.02.626412

**Authors:** Daria Travnikova, Vera Shashkovskaya, Maria Kordyukova, Irina Nosova, Olga Kudryashova, Timofei Zatsepin, Vsevolod Belousov, Tatiana Abakumova

## Abstract

**Abstract:** The oncoprotein Gankyrin/PSMD10 is upregulated in various cancers, including glioblastoma multiforme (GBM), the most difficult tumor to treat. The heterogeneity of this type of tumor requires a multi-targeted approach to overcome radio- and chemoresistance. Here we aim to investigate the effects of Gankyrin inhibition on glioma cell proliferation and its impact on sensitivity to chemotherapy both *in vitro* and *in vivo*. We have employed the RNA interference approach as a valuable tool to downregulate the expression of target genes. Small interfering RNA (siRNA) were formulated into lipid nanoparticles (LNP) to facilitate delivery and overcome challenges, including RNase activity. We demonstrate that knockdown of Gankyrin promotes cell growth arrest of murine (GL261, CT-2А) and human (U87, GBM23, GBM24) glioma cells and induces the generation of reactive oxygen species and apoptosis. Importantly, Gankyrin inhibition enhances the therapeutic effect of temozolomide in murine glioma cell lines and decreases the IC50 of doxorubicin and cisplatin for U87 and GL261 cells. The therapeutic potential of Gankyrin inhibition was shown in patient-derived glioblastoma stem cells, which are responsible for tumor initiation and resistance to therapy. We showed that downregulation of Gankyrin led to decreased cell proliferation *in vitro* and improved survival in xenograft glioma model mice. Our results suggest that harnessing siRNA against Gankyrin may serve as a novel therapeutic strategy for glioblastoma therapy in a multi-drug approach and address challenges in current treatment by sensitizing glioma cells to chemotherapy.

**Highlights:**

- siRNA-mediated knockdown of Gankyrin reduced cell proliferation of murine and human glioma cells, induced ROS generation and promoted apoptosis.
- Inhibition of Gankyrin increased the therapeutic effect of temozolomide in resistant CT- 2A glioma cells and decreased the IC50 concentration of doxorubicin and cisplatin in U87 and GL261 cells.
- Downregulation of Gankyrin inhibited the growth of patient-derived glioma stem cells and increased the survival of NOD/SCID xenograft glioma model mice

## 1. Introduction

Glioblastoma multiforme (GBM) is a WHO grade 4 glioma and the most aggressive type of primary malignant brain tumor that GBM has a low survival rate [1]. Mortality from this disease can reach almost 100% [2] and five-year survival rate is less than 5% [3]. Newly diagnosed GBM is currently treated with surgery followed by radiation and temozolomide (TMZ) chemotherapy [4]. However, the recurrence is about 80% and the major problem of GBMs is chemoresistance, leading to relapses and poor survival [5]. A small number of patients are eligible for repeat surgery or radiation treatment. However, in most cases, further treatment with medications is the only option [6]. Therefore, novel targets and methods for the treatment of GBM, such as immunotherapy, new chemotherapy and/or radiation protocols are studied [7].

Prolonged treatment with TMZ often fails due to the resistance of glioma cells. The heterogeneity of GBM is associated with multiple targets that contribute to glioma progression. So, a multi-targeted approach using the repurposing of various drugs in combination with the inhibition of newly identified targets should be explored. Inhibition of the key pathways involved in chemoresistance could be utilized as a therapeutic strategy to sensitize cancer cells and improve outcomes. Polo-like kinase 1 selective inhibitor, Volasertib, has been shown to be involved in G2/M phase cell cycle arrest and DNA damage leading to apoptosis of glioma cells and preventing tumor growth [8]. Other potential targets for the prevention of chemoresistance in tumor cells are human RecQ helicases, which are involved in DNA repair. It was found that the RecQL4 expression is upregulated in GBM, and its downregulation, followed by delivery of small interfering RNA (siRNA) increases the sensitization of glioma cells to chemotherapy [9]. Bioinformatic analysis revealed that the mitochondrial outer membrane protein (MTCH2) is overexpressed in glioma, and its silencing increased the TMZ sensitivity in human glioma cells, which may serve as a potential target for further research [10].

Gankyrin (also known as PSMD10 or p28^GANK^) is a regulatory subunit of the 19S proteasome and is involved in transporting ubiquitinated proteins to the 20S proteasome for degradation [11]. Multiple human cancers, including hepatocellular carcinoma (HCC), lung cancer, and GBM, have shown dysregulation of Gankyrin [12]. Overexpression of Gankyrin is involved in tumor initiation and progression by regulating several signaling pathways that control cell cycle processes, cell growth, invasion and apoptosis [13]. It has been found that Gankyrin inhibits two tumor suppressor proteins - p53 and the retinoblastoma protein and is also implicated in the modulation of key cellular signaling pathways, including STAT3/AKT, Nf-kB, and mTORC1 [14], [15].

Several investigations showed that inhibition of Gankyrin affect cell growth, and also enhance the chemosensitivity of cancer cells. Previously, we have demonstrated that siRNA-mediated knockdown of Gankyrin led to sensitization of HCC cells to sorafenib [16]. Similar results were demonstrated by D’Souza, A.M with a combination of a small molecule inhibitor and doxorubicin or cisplatin, which led to improved cytotoxicity and apoptosis in liver cancer [17]. A recent study demonstrated the synergistic effect of TMZ and doxorubicin in GBM cells (GBM4, GBM6, and U87), inducing cell apoptosis and potentially serving as a combination therapy for glioma patients [18]. It was also shown that doxorubicin could serve as a first-generation of small molecule inhibitor of Gankyrin [19], so the potency of the combination those of two drugs should be investigated.

A number of studies have investigated the impact of Gankyrin overexpression in GBM tissue on tumor growth and poor prognosis in patients [20], [21]. Yang et al. showed that the inhibition of Gankyrin prevents the proliferation of U251 glioma cells and is involved in cell cycle arrest [13]. Therefore, Gankyrin could potentially be a target and a component of glioma treatment protocols; however, the underlying mechanisms are still unclear and require further investigation [13].

siRNA can serve as both a new treatment approach and a tool for conducting proof-of-concept studies to demonstrate the efficacy of inhibiting specific targets. siRNAs have potential as therapeutic tools for silencing genes that play a major role in the development of malignant glioma phenotypes, such as proliferation, invasion, metastasis, resistance to treatment, and immune escape [22]. However, there are many limitations to the clinical use of siRNAs due to their degradation in the blood, fast elimination, insufficient cellular uptake, blood-brain barrier and immune response [23]. Various systems based on the encapsulation of siRNA into different nanocarriers, such as lipid nanoparticles (LNPs), liposomes, exosomes, polymers and micelles were investigated for delivery to the brain [24], [25]. To date, there are four FDA-approved siRNA drugs, including Patisiran (ONPATTRO^TM^) which is a siRNA agent encapsulated into LNP for delivery to hepatocytes for the treatment of polyneuropathy [26]. Previously, we showed that the siRNA-mediated knockdown of certain targets involved in the N-degron pathway can sensitize tumor cells to chemotherapy in liver cancer [27]. We also demonstrated that siRNA designed for Gankyrin could sufficiently inhibit the mRNA level of Gankyrin in the murine liver after intravenous injection at a low dose (less than 0.1 mg/kg) [12].

In this study, we aim to investigate inhibition of Gankyrin as a novel therapeutic target for GBM treatment utilizing siRNA as a proof-of-concept approach to silence specific genes. Effective knockdown of Gankyrin expression in glioma cells and its delivery to tumor could potentially reduce cell proliferation, inhibit tumor growth and sensitize cells to chemotherapy.

## 2. Results

### 2.1 Gankyrin expression in GBM and efficacy of siRNA-mediated knockdown of Gankyrin in glioma cells

To assess the expression level of Gankyrin/PSMD10 in glioblastoma (GBM) samples and normal tissues, we analyzed publicly available bulk RNA-seq data: the Cancer Genome Atlas (TCGA) for tumor samples and Genotype-Tissue Expression (GTEx) Portal for normal tissues. First, we compared the expression of PSMD10 between tumor samples and healthy brain tissues. We found that PSMD10 was significantly more highly expressed in glioblastoma samples compared to normal brain tissues (median 6.2 vs. 5.8, p-value = 4.1 × 10e^−8^), Fig.1A. Also, we examined the expression of Gankyrin in different healthy tissues. We found that Gankyrin was significantly upregulated in GBM than in all other presented tissues from GTEx Portal (Mann- Whithey test, p < 0.001), while a group of tissues showed notably lower expression (Figure 1B). Interestingly, this group included the heart, liver and pancreas, suggesting that a potential knockdown of Gankyrin in these tissues would not likely result in significant adverse effects.

**Figure 1.**
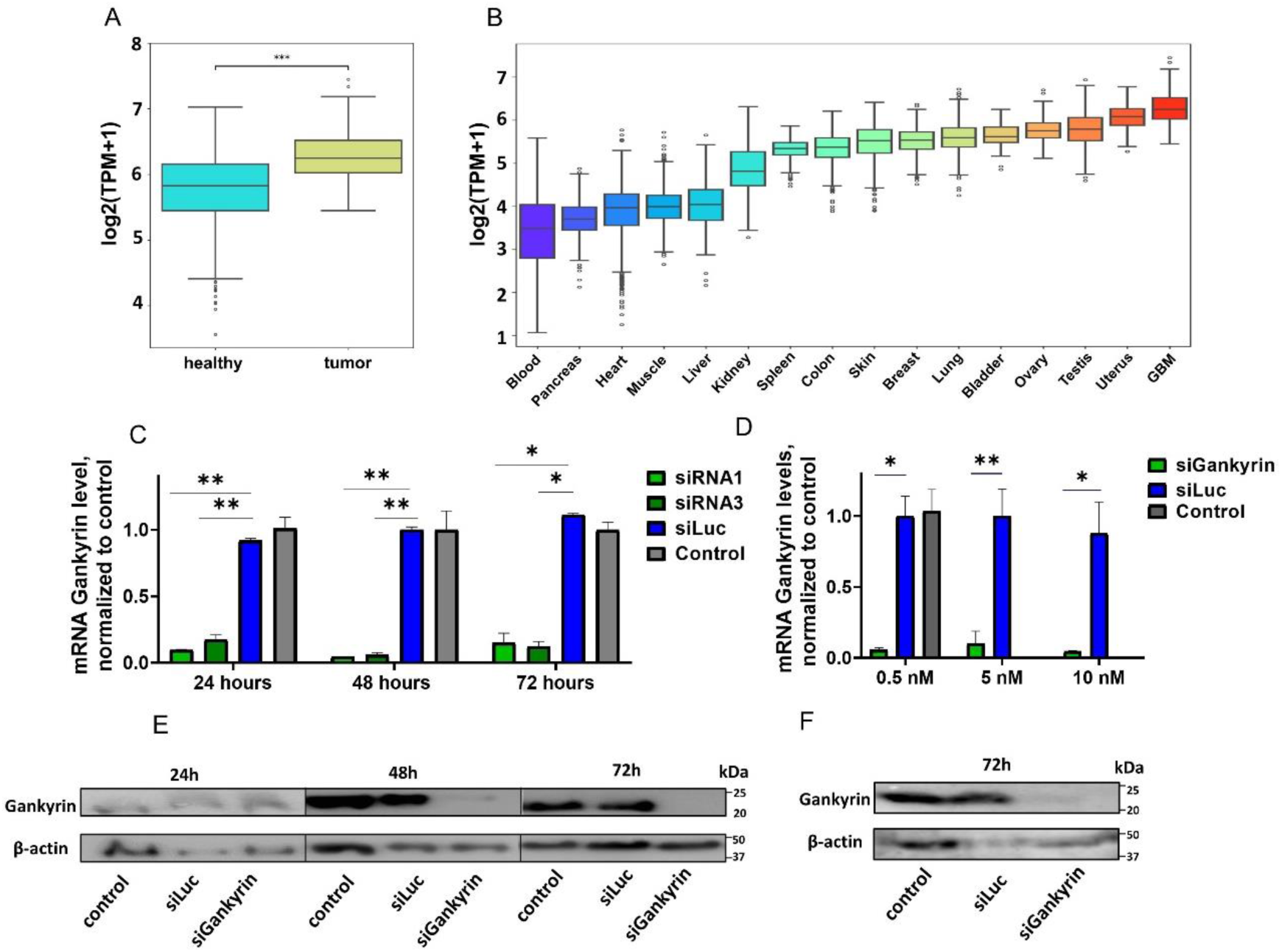
(**А**) Comparison of Gankyrin/PSMD10 gene expression based on RNA-Seq data from human glioblastoma multiforme samples (TCGA) and normal brain tissues (GTEX) (marked as Tumor and Healthy respectively). (Mann-Whitney test, *** p < 0.001). (**B**) Expression profile of Gankyrin in glioblastoma samples (GBM) and different healthy tissues based on RNA-Seq data (C) The effect of siRNA-mediated Gankyrin knockdown in U87 and GL261 cells. (D) mRNA Gankyrin level in U87 cells determined by RT-PCR 24, 48 and 72 hours after transfection with 10 nM of siRNA-1 or siRNA-3. siRNA to firefly luciferase (siLuc) and untreated cells (control) were used as negative controls. (E) mRNA Gankyrin level in GL261 cells in 48 hours after transfection with various siRNA concentrations (0.5 nM, 5 nM, 10 nM). (F) Western blot analysis of Gankyrin protein expression in U87 cells after 24, 48 and 72 h and in GL261 cells in 72 hours after knockdown with siRNA. Three biological replicates were used; results show mean ± SD. (**p<0.01, *p<0.05, multiple t-test).

To achieve specific suppression of murine and human Gankyrin (PSMD10) mRNA, three siRNA, which were designed earlier [12], [16], were tested in GL261 and U87 glioma cells (siRNA-1, siRNA-3, siRNA-6). The most efficient siRNA for Gankyrin inhibition were selected and utilized in subsequent investigations (siRNA-1 and siRNA-3). We demonstrated that both siRNAs can effectively knock down Gankyrin in U87 glioma cells. An siRNA concentration of 10 nM effectively reduced mRNA by 90% after 24 hours of incubation (Fig.1С). It also resulted in decreased protein levels in 72 hours after transfection in all cell lines (Fig. 1E-F).

### 2.2 Gankyrin knockdown affects glioma cell proliferation, induces apoptosis, cell cycle, reactive oxygen species (ROS)

Previously, we determined that 72 hours the downregulation of Gankyrin lead to more than 90% reduction of mRNA (RT-PCR) and protein (Western blot). We evaluated cell metabolic function after transfection with Alamar Blue assay [28] varying the siRNA concentrations (1 nM, 5 nM, 10 nM) and the exposure time from 48 hours to 96 hours. Murine GL261 glioma cell proliferation was most significantly inhibited within a 96-hour transfection period. Cell growth was suppressed by at least 40% by adding 5 nM siRNA and more than 50-60% with 10 nM siRNA (Fig.2A-B). This cell line is extremely sensitive to Gankyrin as cellular viability experiences a substantial decline even following a 72-hour transfection period with a concentration as low as 1 nM of siRNA (Fig. S1C). Meanwhile, human cell line U87 is more resistant to Gankyrin knockdown, decreasing survival rate by up to 10% and 20% with 5 nM and 10 nM siGankyrin, respectively (Fig.2C-D). CT-2А has shown no significant difference in viability after knockdown (Fig.2E-F) at 96 hours. We have also analyzed the knockdown effects in mouse astrocytes. Our findings demonstrate that cell viability remains unaltered with a concentration of 10 nM of siRNA for 48 hours, as confirmed by microscopic imaging data (Fig.S2).

**Figure 2.**
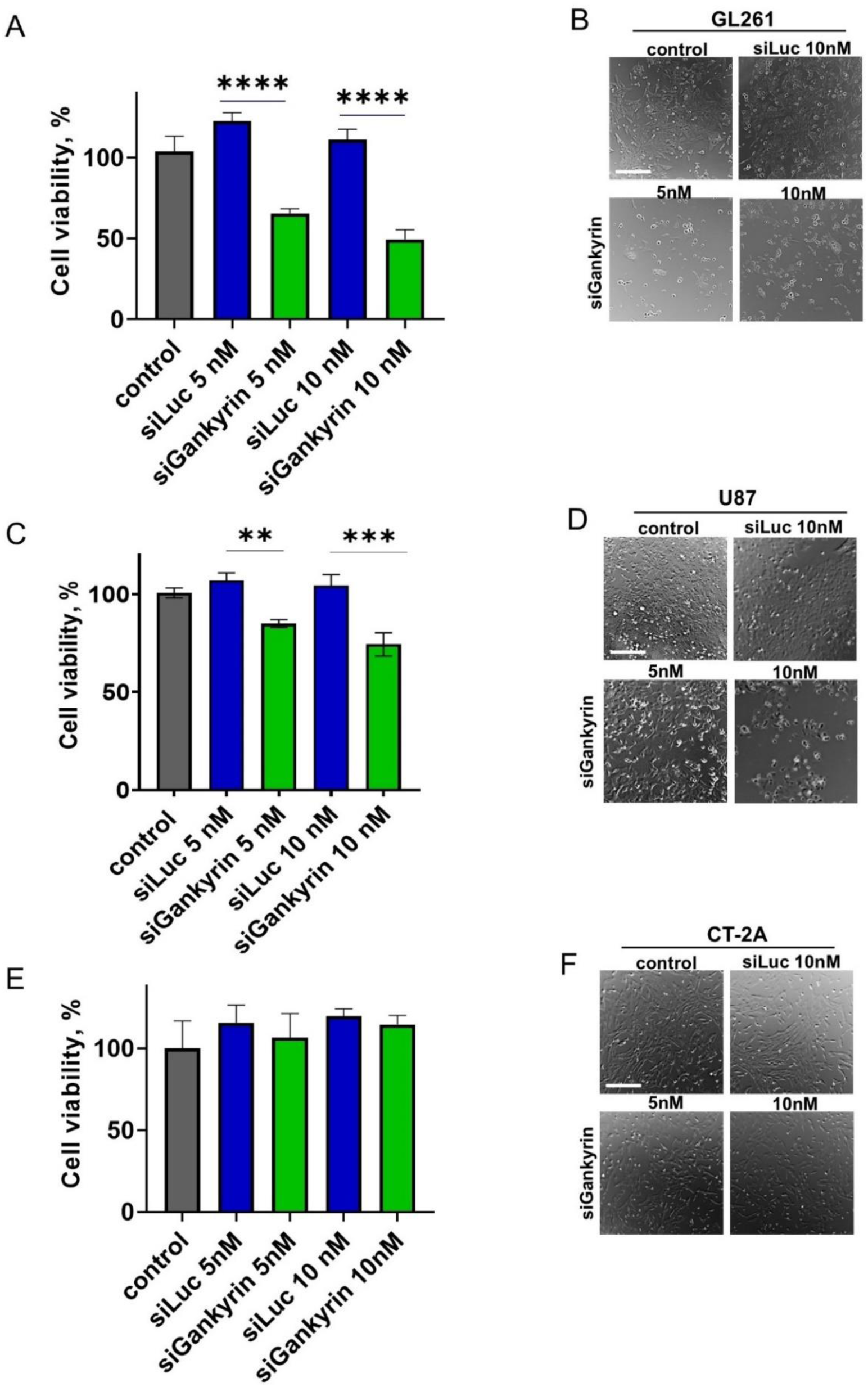
Cell viability after transfection with siRNA Gankyrin normalized to untreated cells (control) (A) GL261 (C) U87 and (E) CT-2A viability in 96 hours after transfection with siRNA to Gankyrin in concentrations of 5 nM and 10 nM compared to the control siRNA (siLuc). (B, D, F, H) Images of the untreated cells and cells treated with siRNA cells (GL261, U87, CT-2А) 48 hours after transfection. Scale bar 50 µm. Five biological replicates per sample were used; results show mean ± SD. (** p < 0.01,*** p < 0.001, **** p < 0.001, One-way ANOVA)

In addition, we investigated the role of oxidative stress as a critical factor in cell death, in particularly accumulation of reactive oxygen species (ROS). Our findings indicate that fluorescence intensity in siRNA Gankyrin-treated cells was significantly elevated across all cell lines when compared to internal controls as detected by H2DCFDA assay (Fig. 3A). This substantial increase in fluorescence suggests that inhibition of Gankyrin leads to a marked generation of ROS, which may subsequently result in cell death and apoptosis.

**Figure 3.**
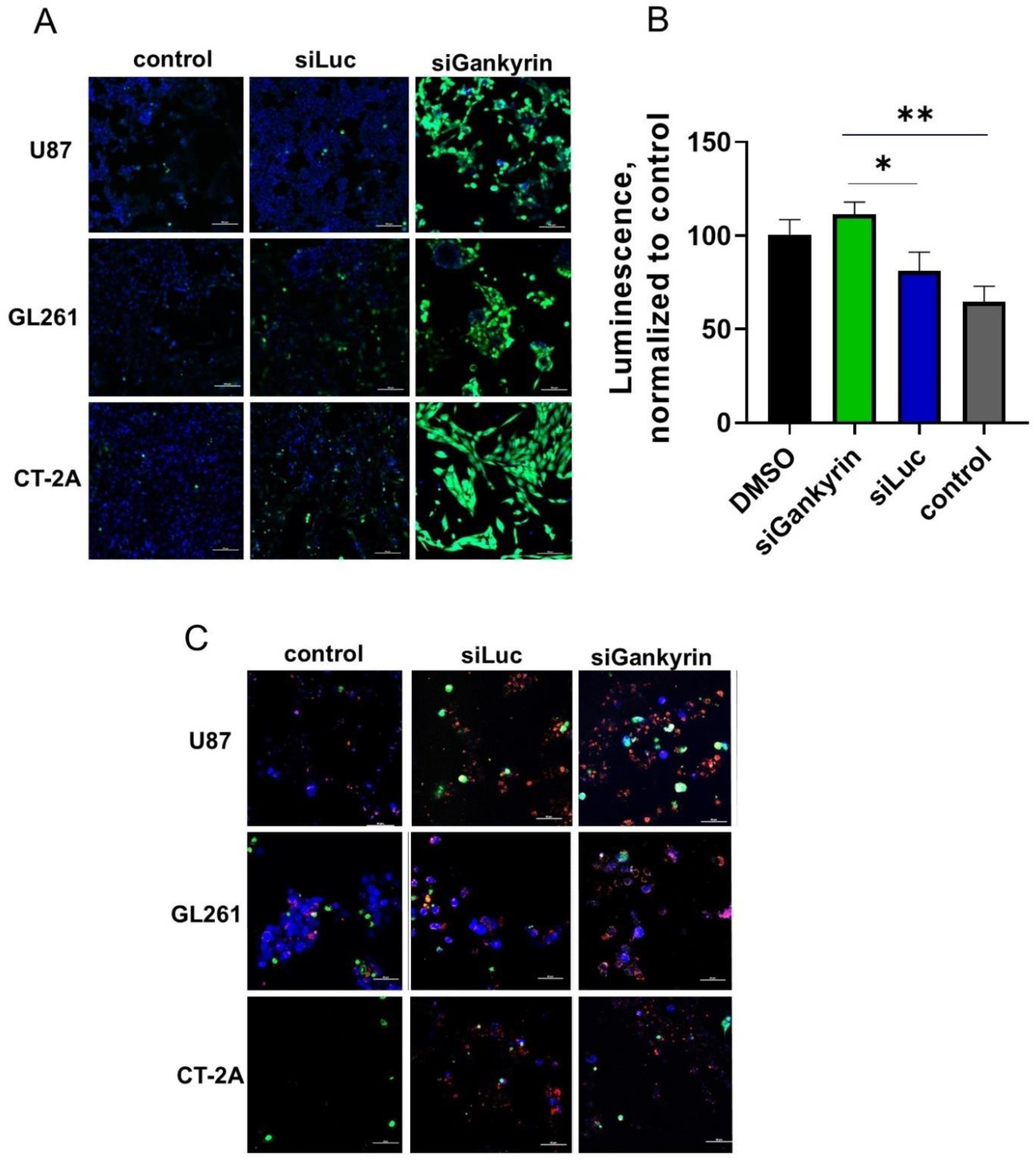
Inhibition of Gankyrin can promote apoptosis and increase ROS generation. (A) Analysis of ROS generation with H2DCFDA assay in 24 hours after transfection with siRNA to Gankyrin (10 nM). Scale bar 100 µm. (B) Caspase-Glo® 3/7 Assay: normalized luminescence signal from apoptotic U87cells, 48h-transfection, 10 nM. Cells treated with 1% DMSO were used as a group for normalization. (C) Analysis of cell death by the apoptosis/necrosis assay 48 hours after transfection with siRNA to Gankyrin (10 nM). The live cells are indicated by blue, the apoptotic cells by red, and the necrotic cells by green. Scale bar 50 µm. Four biological replicates per sample were used; results show mean ± SD. (**p<0.01, *p<0.05 ns - non-significant, one-way ANOVA).

The mechanism of cell death after Gankyrin knockdown was evaluated using an Apoptosis/necrosis staining kit with confocal microscopy. We assume that a significant number of the cells in all lines (U87, GL261, CT-2А) follows a pathway that initiates apoptosis after Gankyrin knockdown (Fig.3). However, it should be noted that prolonged incubation with control siRNA was also accompanied by the presence of apoptotic cells.

To further confirm the impact of siRNA Gankyrin on the induction of apoptosis in glioma cells, we assessed caspase-3/7 activity as a measure of apoptotic signaling. Inhibition of Gankyrin led to a 30-37% increase in apoptotic activity compared to the control group (Fig.3B). This finding suggests that Gankyrin knockdown may promote apoptosis, albeit without a corresponding increase in the total number of apoptotic cells detected in the initial analysis. Notably, we also detected an increase in the levels of caspase-1 through Western blot analysis following Gankyrin knockdown (Fig. S1D). Caspase-1 is a crucial enzyme involved in inflammatory responses and is primarily known for its role in the cleavage of pro-inflammatory cytokines such as IL-1β and IL-18. The fact that its levels rose after Gankyrin knockdown suggests a potential regulatory relationship between Gankyrin and the inflammasome pathway.

To study the effects of Gankyrin downregulation on the cell cycle, we incubated glioma cells for 48 hours to avoid significant cell death. We analyzed the cell cycle with flow cytometry after propidium iodide staining (Fig.4). Downregulation of Gankyrin resulted in an increase in the cell number in the G1 phase and a decrease in the S/G2 phases in the U87 cells. In siGankyrin-treated U87 cells, 78±2% of the cells were in the G1 phase, compared to 66– 67% in all control cells (Fig. 4). In the GL261 cell line, 70% of siRNA-treated cells were in the G1 phase, compared with 47% in control (PBS) cells, while the difference in the S/G2 phase was even more remarkable: 18% for siGankyrin, 31% for siLuc, and 63% for control. In the CT-2А cells, we did not observe any significant differences between cell numbers in phases for transfected and control cells. We believe that further comprehensive analysis of cell lines with varying sensitivity to siGankyrin could reveal the biomarkers that could be used to explain these findings and translate them to clinical practice.

**Figure 4.**
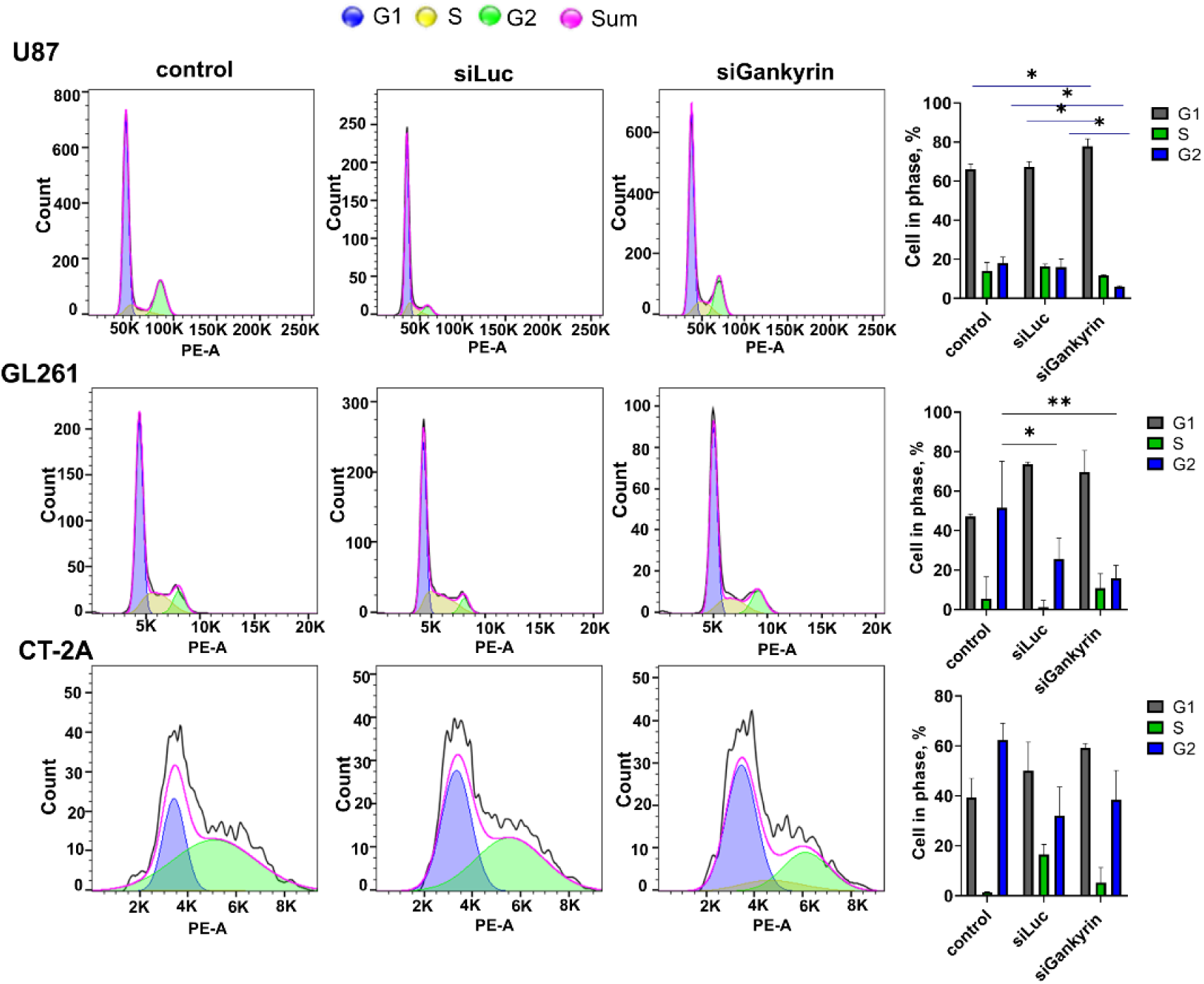
Inhibition of Gankyrin by siRNA induces G0/G1 arrest in glioma cells. Cell-cycle distribution in different phases was assessed by flow cytometry and FlowJo calculation. Transfection was prepared for 48 hours with 10 nM siGankyrin in U87, GL261 and CT-2А. siRNA against firefly luciferase (siLuc) was used as a negative control. Three biological replicates were used; results show mean ± SD. (*p<0.05, ** p < 0.01, multiple comparison-test ANOVA was used)

### 2.3 Downregulation of Gankyrin sensitizes glioma cells to chemotherapy (doxorubicin, cisplatin and TMZ)

The ability of siRNA to Gankyrin to sensitize liver cancer cells to the chemotherapeutic drug sorafenib has previously been demonstrated [16]. In this research we have also hypothesized that it might not only initiate cell growth arrest, but also sensitize cancer cells to chemotherapeutic drugs such as TMZ, doxorubicin and cisplatin. We investigated the effect of chemotherapeutic drugs on glioma cell lines with different sensitivity (Table S1) after siRNA-mediated knockdown and analyzed cell viability using Alamar blue assay and phase-contrast microscopy.

As TMZ is a standard treatment for GBM, we sought to enhance its therapeutic effect by inhibiting Gankyrin in cell lines with different TMZ-sensitivity: the IC50 values for CT-2А and U87 exceeded 250 µM, while GL261 cells were sensitive to TMZ (IC50=21 µM). We inhibited the expression of Gankyrin with siRNA for 48 hours in all cell lines and then treated them with different doses of TMZ (0-2000 µM) for 72 hours. Here, we demonstrated that at lower doses, TMZ in combination with siRNA effectively inhibited the growth of CT-2А cells and made them more susceptible to TMZ. The cells exhibited a response profile comparable to that observed in TMZ-sensitive GL261 cells. TMZ alone at concentration 250 µM reduced cell survival by 15%, but when CT-2А were pre-treated with siRNA against Gankyrin, cell mortality was 79% (or a survival rate of 21%) (Fig.5A). The survival rate of GL261 cells was approximately 22% at the same concentration of TMZ without knockdown (Fig.5B). This signifies that CT-2А cells after knockdown start to respond to TMZ in the same way as sensitive cells. Therefore, Gankyrin knockdown sensitized CT-2А to TMZ chemotherapy, causing their to respond like non-resistant cells.

**Figure 5.**
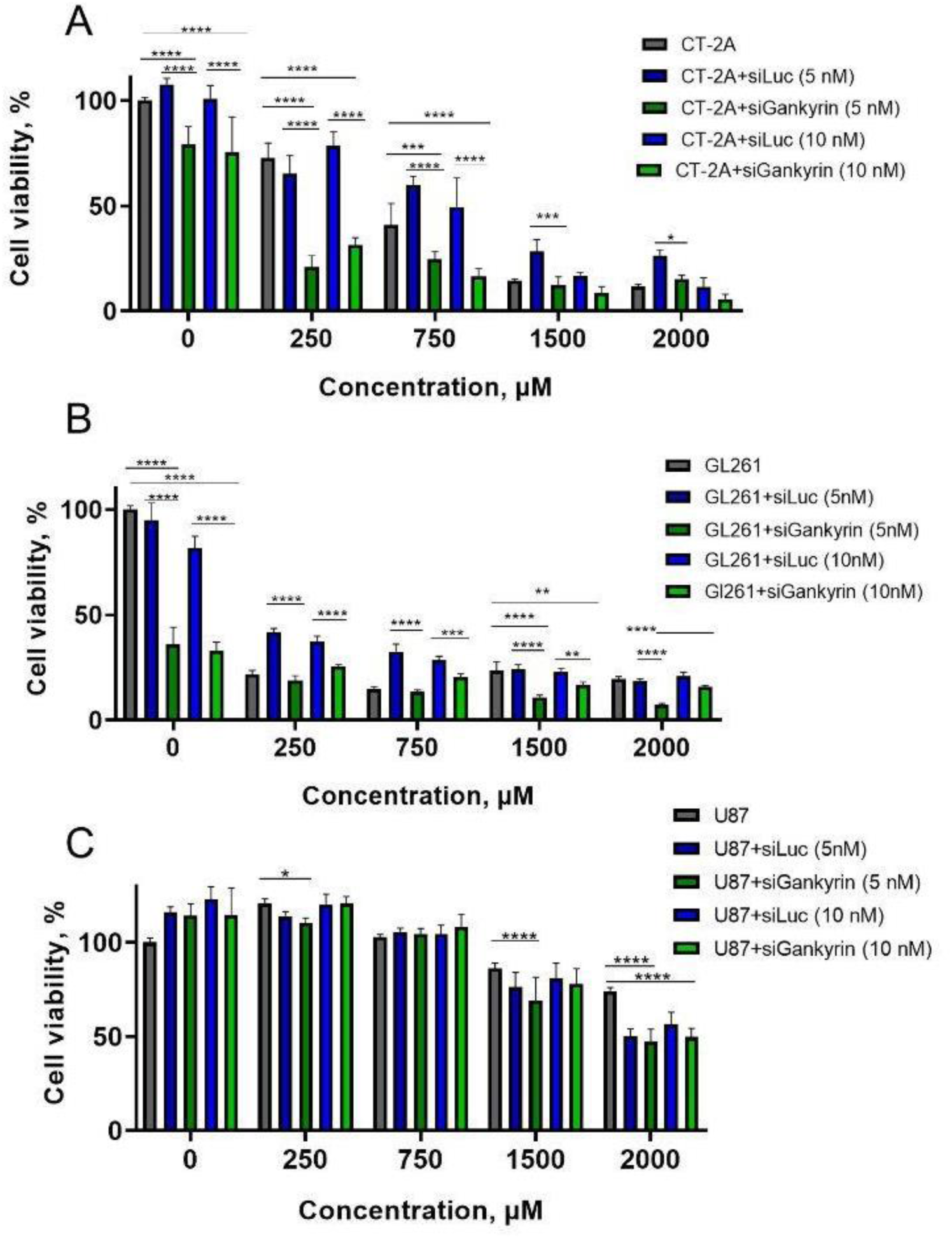
Therapeutic synergy between siRNA to Gankyrin (5 nM, 10 nM) and TMZ at GL261 (A), CT-2А (B) and U87 (C). Transfection lasted for 48 hours followed by a 72-h incubation of the drug. SiRNA against firefly luciferase (siLuc) was used as negative control. Six biological replicates were used; the results show mean ± SD. (* p<0.05, ** p < 0.01, *** p<0.001, **** p< 0.0001, multiple comparison-test ANOVA was used)

Furthermore, studies were conducted with other drugs, such as cis and dox, given their potential efficacy as second-line drugs for GBM. The cells were pretreated with siRNA for 2 days to inhibit Gankyrin and then incubated for 24 hours. As previously, GL261 cell also demonstrated a notable response to 5 nM siGankyrin alone, resulting in an 82% survival rate. When treated with cis and dox alone, GL261 cell viability was 82% and 72% respectively. The addition of cytostatic drugs after siRNA treatment showed an additional reduction in cell viability of up to 20% (overall 43-46% increase in cytotoxicity compared to control). A similar trend was observed in cytotoxicity experiments conducted with the U87 cells. We have observed that Gankyrin knockdown enhances the efficacy of drug therapy by up to 28%–32%. In contrast, CT-2А cells were less sensitive to Gankyrin inhibition, with no observable cell death following knockdown (Fig. 2E, 6E), which confirms previous results. Nevertheless, Gankyrin knockdown tends to enhance the response to cis therapy, with a decline in viability from 78% (siLuc+cis) to 53% (siGankyrin+cis). (Fig.6 (E-F)).

**Figure 6.**
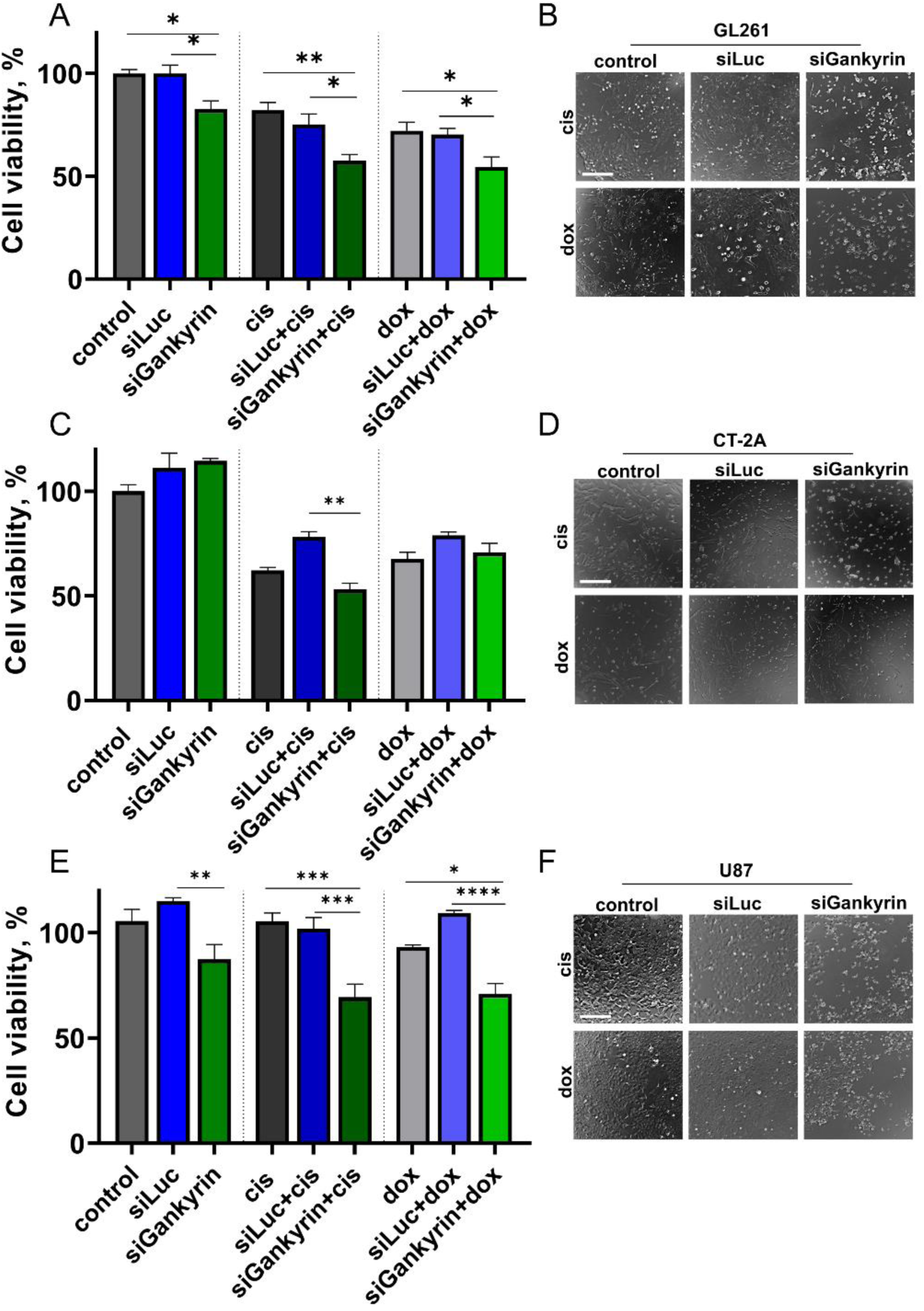
Therapeutic synergy between siRNA Gankyrin and cis, dox. Transfection for 48 hours followed by 24 hours of drug incubation. (A, B) GL261 viability at 5 nM siRNA incubation. (C, D) CT-2А viability at 10 nM siRNA incubation. (E, F) U87 viability at 10 nM siRNA incubation. siRNA to firefly luciferase were used as negative control. Scale bar 100 µm. Five biological replicates were used; the results show mean ± SD. (***p<0.001, **p<0.01, *p<0.05 ns - non-significant, one-way ANOVA).

### 2.4. Biodistribution and efficacy of siGankyrin – LNP delivery after intratumoral and intravenous injection to orthotopic glioma model mice

Although the downregulation of Gankyrin in vitro has been successfully achieved and its therapeutic potential has been demonstrated, the difficulties associated with delivery to the tumor site and knockdown in tumor *in vivo* remains challenging. Previously, we have shown that intravenous injections of siGankyrin -LNP led to downregulation of mRNA in liver tissue. Here, we compared different delivery routes to investigate the optimal route for further proof-of-concept experiments on the therapeutic potential of siGankyrin for the glioma treatment.

First, we validated the tumor growth of glioma model GL261 - the most sensitive to Gankyrin inhibition and chose 7 and 21 days after implantation as the time point for early-stage and late-stage glioma, respectively. We analyzed the biodistribution of siRNA-LNP after intravenous (1 mg/kg) and intratumoral (1 µg) injection by: 1) fluorescent in vivo imaging of glioma-bearing mice after different routes of injection and 2) analysis of the efficacy of knockdown of the target gene (Gankyrin) in glioma and healthy brain hemispheres 48 hours after injection.

Efficient siRNA delivery by intravenous administration was problematic at the early stages, as the IVIS images represent no fluorescent signal in the perfused brain (Fig. 7A). RT-PCR data also supported these findings, showing that there was no significant difference between brain tissue from the control group and glioma after siRNA-LNP injection (Fig. 7B). Intratumoral injection was preferable for siRNA-LNP for this stage and allowed effective downregulation of Gankyrin (Fig. 7D). In contrast, intravenous injection of siRNA-LNP at the late-stage glioma led to significant accumulation of nanoparticles in the tumor site as confirmed by IVIS images (Fig. 7E). Moreover, it was demonstrated that untreated gliomas (with siLuc injection) exhibited the highest levels of Gankyrin expression, while treated gliomas showed a significant reduction in mRNA Gankyrin level after siRNA-mediated knockdown (Fig. 7F). Confocal microscopy confirmed these findings, explaining revealing significant disruption of the blood-brain barrier and demonstrating the efficient targeting and knockdown of genes into tumors by siRNA-LNP at late stages of glioma.

**Figure 7.**
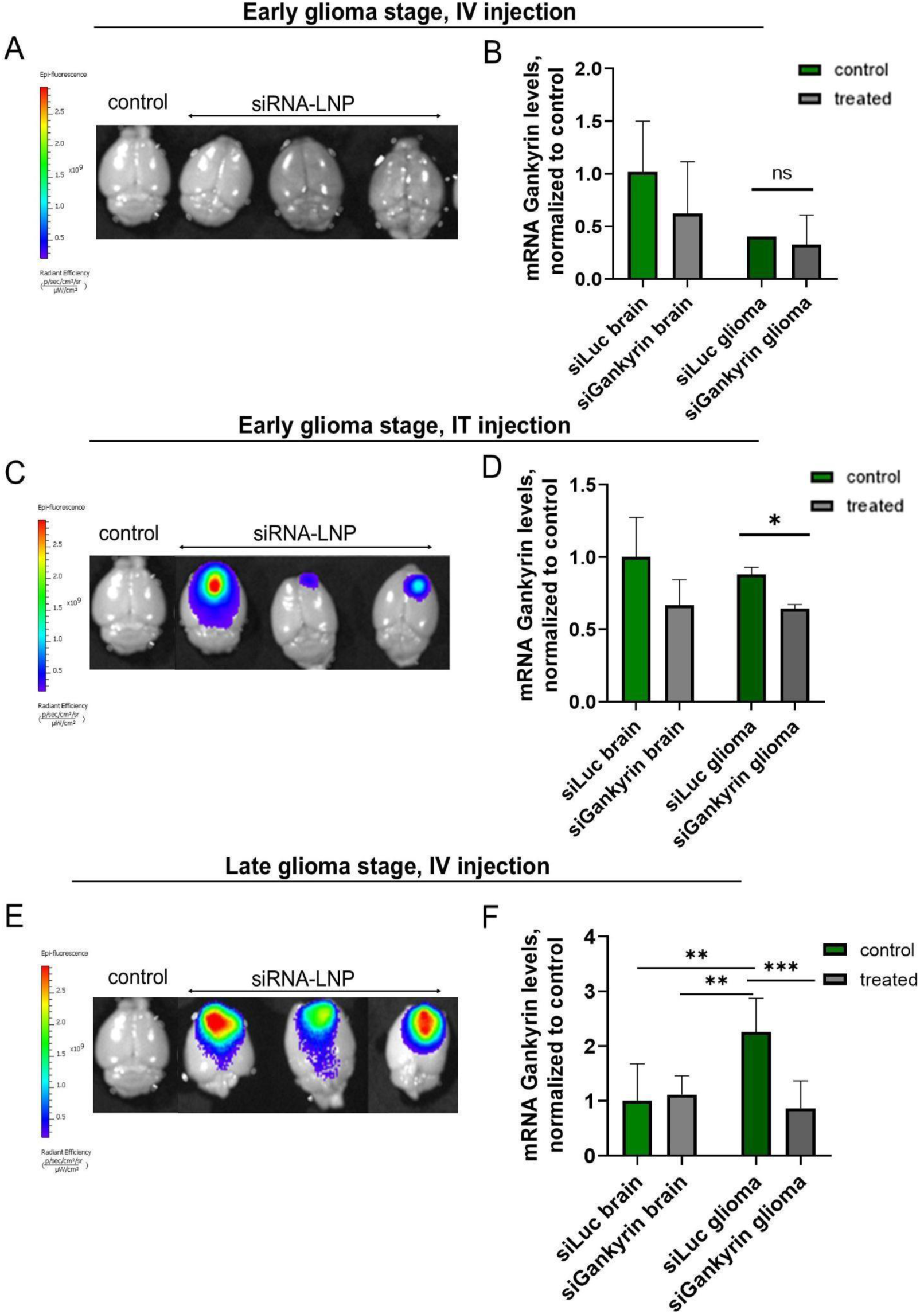
Biodistribution of siRNA-LNP-cy5 at 7 days (early-stage glioma) and 21 days (late stage glioma) after implantation of GL261 cells after intravenous (A, E) and intratumoral injections (C). Mice injected with PBS were used as the negative control (B, D, F) mRNA Gankyrin level in glioma and healthy hemisphere (brain) samples after different routes of injection. Five biological replicates were used; the results show mean ± SD. (***p<0.001, **p<0.01, *p<0.05 ns - non-significant, one-way ANOVA).

### 2.5 siGankyrin effects on patient-derived GBM in vitro and in vivo and perspectives of translation

Chemoresistance and aggressiveness of GBM cells is a crucial problem and significantly affects treatment outcomes, leading to poor survival and further tumor growth. Recently, it was shown that Gankyrin is overexpressed in GBM tissue [29]. We went further and analyzed mRNA Gankyrin level in patient-derived glioma stem cells and chose two glioma lines with different expressions of Gankyrin - GBM23 and GBM24 (Fig. 8A).

**Figure 8.**
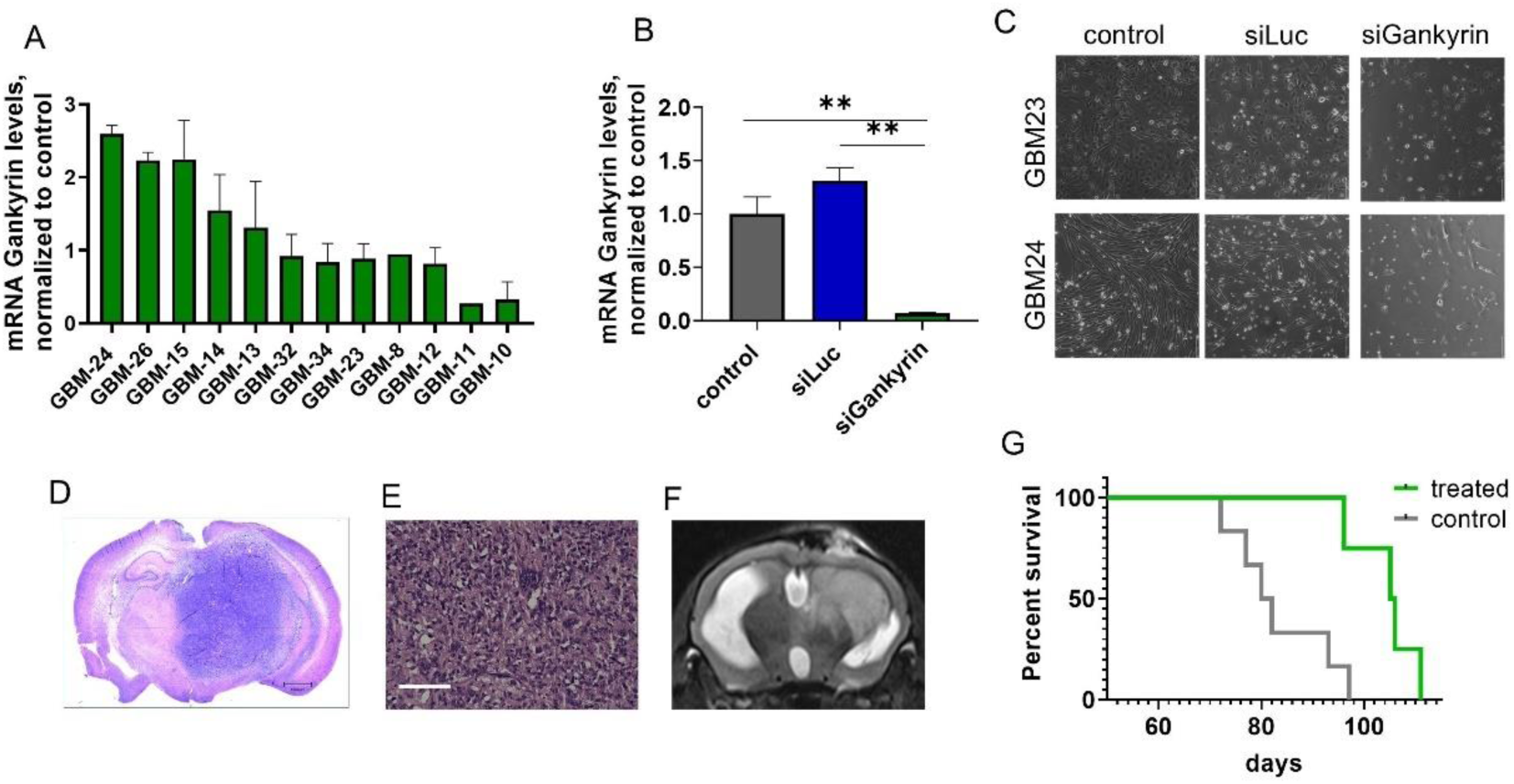
siGankyrin effects on patient-derived GBM in vitro and in vivo. (A) mRNA Gankyrin level in patient-derived glioma stem cells. (B) mRNA Gankyrin level in GBM23 after knockdown by RT-PCR. Transfection was performed 2 times every 72 hours at 10 nM concentration. Multiple t-test was used (**p<0.01) (С) GBM23 and GBM26 viability at 10 nM siRNA incubation for 48 hours. (D, E) Histological analysis from xenograft patient-derived glioma model (GBM23), hematoxylin eosin staining. Scale bar 1000 μm and 120 μm. (F) T2- weighted MRI image from NOD/SCID glioma model with GBM23 at 3 month after implantation. (G) Survival of mice with implanted GBM23 cells. Log-rank test was used (p<0,01).

First, we were able to successfully transfect primary cells and downregulate human Gankyrin by 93% in the selected patient-derived GBM lines (Fig. 8B). Next, we confirmed the previous results with immortalized glioma cells (Fig. 2 (A-B, E-F)) and demonstrated that siGankyrin transfection led to inhibition of cell growth and death (Fig. 8B). Interestingly, the lethality of the treatment was more pronounced in GBM cell lines than in the human U87 cells.

Finally, we established and validated the growth of the xenograft glioma model with patient-derived GBM23 cells implanted into the striatum of NOD/SCID mice. GBM23 cells exhibited aggressive growth patterns *in vivo,* characterized by rapid expansion and invasiveness, which is typical of high-grade gliomas and was confirmed by histological staining (Fig. 8 D, E). MRI scans demonstrated significant increases in tumor volume and supraventricular growth (Fig. 8F). siGankyrin was administered intratumorally at 72-hour intervals (4 times, dose 1 μg siRNA) 1 month after tumor initiation. The survival analysis indicated a significant difference in median survival times: 106 days for the siRNA Gankyrin-treated mice, compared to 81 days for the control group (Fig.8D), indicating the therapeutic potential of targeting Gankyrin in the treatment of GBM.

## 3. Discussion

GBM is one of the most aggressive and malignant types of cancer in humans with an urgent need for new treatment options, including novel targets and anticancer drugs. Chen A. observed that all members of the PSMD family showed higher protein expression levels in GBM than in normal brain tissue [13]. In our study, we investigated the role of PSMD10 (Gankyrin) in the GBM cell proliferation alone and its efficacy in sensitizing cells to chemotherapy. It was highlighted that the GBM cells become resistant to the chemotherapeutic agents due to the high expression of PSMD10 [21]. Additionally, it was shown that Gankyrin could serve as a potent prognostic factor for estimation of the overall survival of patients with GBM [21]. In our study, we established that Gankyrin is expressed differently in patient-derived GBM stem cells. However, the mRNA Gankyrin level did not correlate with half-inhibitory concentration of TMZ, so other possible biomarkers remain to be discovered.

The use of siRNAs has emerged as a promising strategy for inhibiting specific gene expression, including the oncogene Gankyrin, in cancer research [14]. Our siRNAs demonstrated rapid reduction of mRNA and protein levels, proving its high efficiency in targeting Gankyrin in immortalized and importantly in primary GBM cell cultures, a challenging task [30].

After establishing the successful conditions of knockdown, we evaluated the impact of Gankyrin knockdown on glioma cell viability and proliferation. Interestingly, we observed a decrease in the survival rate of certain glioma cell lines (GL261, GBM23, GBM24) after knockdown, suggesting that Gankyrin may play a more important role in the metabolism of these cells than those less sensitive to its inhibition. Among the cell lines tested, GL261 showed the highest sensitivity to Gankyrin inhibition, which could be attributed to p53 mutations resulting in increased c-Myc expression. Several previous studies have linked Gankyrin to p53 degradation [31].

Further, we demonstrated that the knockdown of Gankyrin expression induced ROS generation even in the CT-2А cell, which appears to be insensitive to siRNA alone in the cell viability test. This phenomenon has been linked to previous findings in studies of PSMD10 involvement in other cancers or ROS generation by other PSMD family proteins [21]. The mechanism likely involves modulation of the ROS via the NF-κB pathway [32]. In a study by J. Wang et al., it was observed that decreasing Gankyrin levels triggered ROS production, which initiated the activation of p38 [32]. The subsequent interaction of p38 with Bax could potentially disrupt the mitochondrial transmembrane potential, causing the release of cytochrome-C from the mitochondria into the cytosol. This cascade may lead to the activation of caspase-9, ultimately initiating the mitochondrial pathway of apoptosis. Similar effects were observed in U87 and GL261 cells, where apoptosis occurred following Gankyrin knockdown, although the difference between siGankyrin and siLuc treatments is only approximate.

Our study elucidated a significant relationship between Gankyrin downregulation and cell arrest at the G1/S phase in the U87 and GL261 cell lines, suggesting a potential role for Gankyrin maintaining the balance between cell cycle and apoptosis [33]. It has been indicated that the high expression of Gankyrin is associated with the degradation of RB1 and TP53, the main regulators of cell growth [34]. Furthermore, it was revealed that it possibly happened through Gankyrin interactions with other key cell cycle regulators such as CDK4 and p21, p27, which may have implications for therapeutic strategies targeting cell proliferation in GBM [35][34].

Chemoresistance is a major obstacle in the treatment of glioma, leading to poor patient outcomes despite advances in therapy. The genetic diversity of tumor cell populations leads to a wide range of target genes involved in resistance. To overcome this challenge, multi-drug protocols such as CUSP9 (now CUSP9v3) have been utilized [36], and the search for new targets is still ongoing to avoid cancer cells becoming insensitive to chemotherapy. In our study, we used three cell lines (CT-2A, GL261, U87) with different sensitivity to common chemotherapeutic drugs, such as TMZ, dox and cis. We have shown that Gankyrin knockdown by siRNA sensitizes resistant cells to TMZ, causing their response to mirror that of non-resistant cells. This phenomenon was demonstrated in murine cells (GL261, CT-2A) and, to a lesser extent, in human U87 cells. The molecular mechanism underlying cell sensitization to TMZ under Gankyrin inhibition has not been thoroughly investigated, although some studies have suggested that PSMD10 expression increases in U87 cells following TMZ exposure [37][35]. However, Zheng et.al. demonstrated the synergistic therapeutic effect in 48 hours after co-administration of siRNA to retinoblastoma binding protein 4 and TMZ in U87MG and U251MG cells [38]. It was also shown that the combination of siRNAs to epidermal growth factor receptor and galectin-1 increased the sensitivity of U87 cells to TMZ and induced cell death [39]. We have also investigated the standard cytostatic drugs such as dox and cis and found that inhibition of Gankyrin improved the efficacy of chemotherapy by 20%. Targeting Gankyrin holds potential for improving treatment outcomes, potentially in synergistic regimens or as a pre-treatment.

The delivery of siRNA to the brain is a challenging task, however, some researchers have raised concerns regarding their potential application in non-liver tumors [40]. The presence of the blood-brain barrier interferes with the penetration of siRNA to the tumor site, but there are several approaches to facilitate delivery: 1) surface modifications of nanoparticles with various ligands and receptors [22]; 2) microbubble-assisted delivery using focused ultrasound [41] 3) convection-enhanced delivery [25]. Recently, a significant number of clinical trials have been focused on the use of intratumoral delivery, especially in the resection cavities left after stereotactic biopsy procedures [42] including oncolytic viruses [43], [44]. siRNA can be administered directly into the tumor, or the other ways for Gankyrin inhibition could be explored, such as short hairpin RNA (shRNA) via viral vectors or small molecule inhibitors. For instance, shRNA Gankyrin repressed cell growth, motility, invasiveness and tumor formation in vivo in esophageal squamous cell carcinoma [45], while small molecule cjoc42 effectively bind to Gankyrin and improved sensitivity to chemotherapy in hepatoblastoma cells [17].

In our study we have demonstrated that successful siRNA-mediated knockdown could be achieved either via intratumoral injection at early-stage glioma or intravenous injection at later stages. Intratumoral injections could be a feasible delivery method for siRNA-LNP with its own advantages as a part of convention-enhanced therapy. Local administration of siRNA-LNP can help reduce systemic exposure to the drug, thereby potentially minimizing systemic side effects and improving the safety profile of the treatment. We demonstrated that the liver showed the highest accumulation of siRNA-LNP after intravenous injection (Fig. S3). However, we do not expect any tissue damage as the expression of Gankyrin is close to negligible in normal organs [17], [46]. Comparable safety findings have been reposted in studies of Gankyrin in various other types of cancer [16].

## 4. Conclusions

We selected the most effective siRNAs against Gankyrin that downregulate its mRNA and protein expression up to 90% in glioma cell lines. The knockdown of the Gankyrin/PSMD10 gene decreased cell proliferation, induced ROS generation, and led to apoptosis of all used glioma cell lines. Downregulation of Gankyrin with siRNA increased temozolomide sensitivity in resistant glioma cells (СT-2A), aligning them with non-resistant cells. Moreover, Gankyrin inhibition was also observed to have a synergistic effect with doxorubicin and cisplatin increasing its cytotoxicity. We have also successfully demonstrated that the downregulation of Gankyrin affects cell growth and survival in patient-derived glioma stem cells. This improved the survival rate of NOD/SCID mice with the xenograft glioma model, making Gankyrin a promising target for further research into improved glioblastoma treatment.

## Acknowledgments

We would like to thank Tatiana Prikazchikova for formulation of LNP for biodistribution and in vivo studies and Olga Patsap for histological staining.The MRI and body fluorescence were performed using the equipment of the Core Facility of Pirogov Russian National Research Medical University.

This project was supported by the grant Russian Science Foundation 22-75-10151 and the grant of Ministry of Health 124021400005-3

## SUPPLEMENTARY

### Materials and methods

#### Cell culture

GL261 and CT-2A murine glioma cell lines were kindly provided by Dr. Aleksei Stepanenko from the Department of Fundamental and Applied Neurobiology of V. P. Serbsky Federal Medical Research Center of Psychiatry and Narcology. U87 cell line was obtained from ATCC Cells and cultured in Dubecco’s Modified Eagle Medium (DMEM) (HiMedia, India) supplemented with 10% fetal bovine serum (FBS) (Serva, Germany), 1% streptomycin-penicillin (PanEco, Russia) and 2 mM L-glutamine (Gibco, USA) at 37◦C with 5% CO2.

#### Real time qPCR (RT-PCR)

Total RNA was isolated using the ExtractRNA reagent (Evrogen, Russia), followed by precipitation with isopropanol, according to the manufacturer’s instructions. 1 μg RNA was used for cDNA synthesis using the Maxima cDNA reverse transcription kit (Thermo Fisher Scientific, USA) or MMLV kit (Evrogen, Russia). Quantitative Polymerase chain reaction was performed using LightCycler 96 (Roche, Switzerland) equipment and qPCR mix-HS SYBR master mix containing SYBR Green I dye (Evrogen, Russia). The primer sequences are provided in Table 1. The mRNA expression of the genes of interest were quantified using the 2-ΔΔCt method, GAPDH was used as a housekeeping gene. The measurements were carried out in three technical replicates for each gene.

**Table 1.**
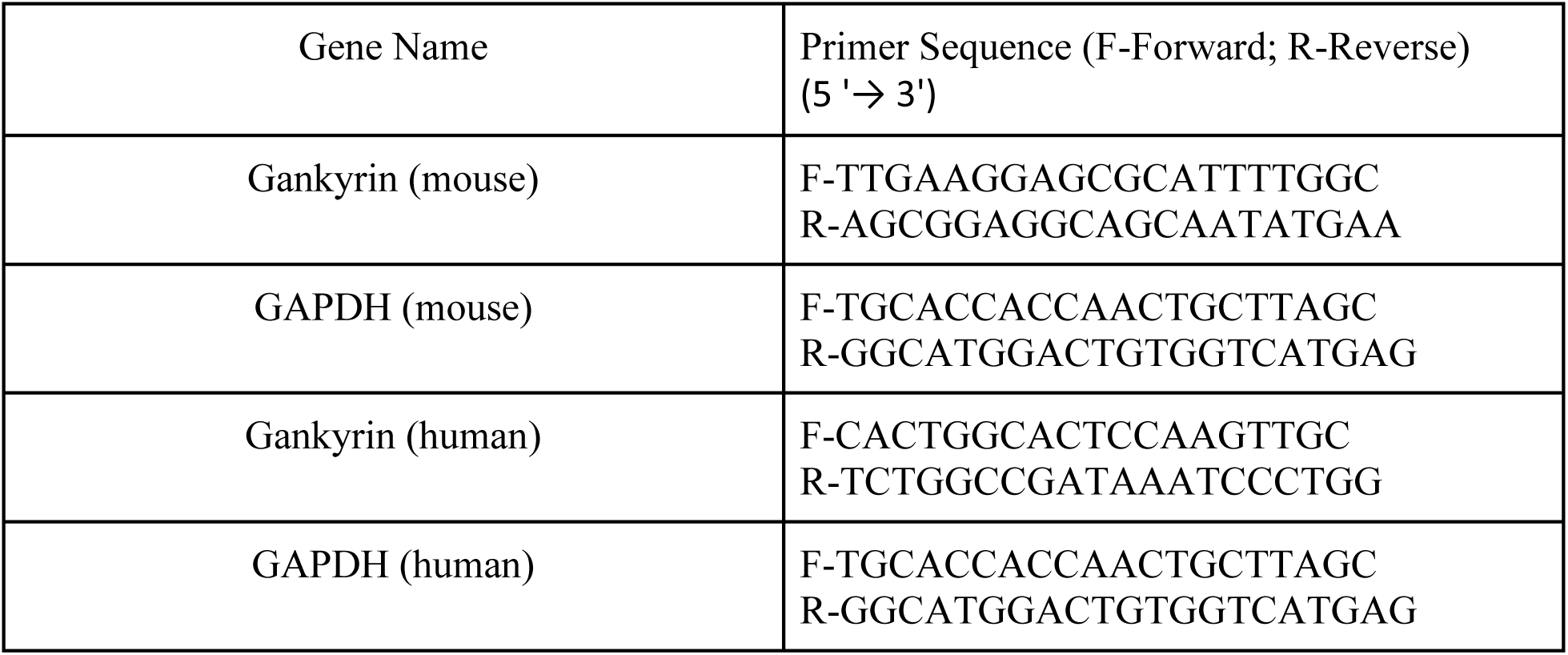
The primer sequences used in RT-PCR.

**Table 2.**
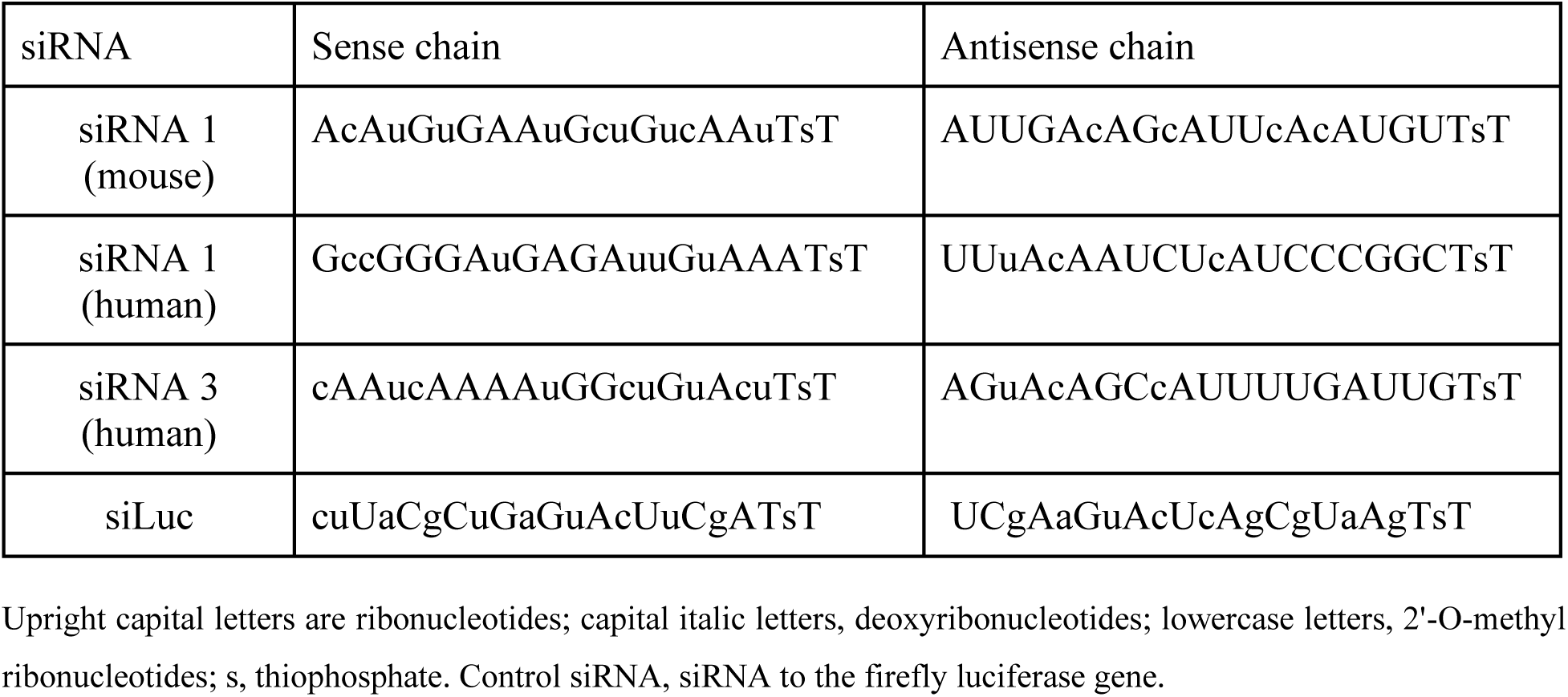
siRNA nucleotide sequences [12], [16].

#### RNA sequencing analysis

We used publicly available bulk RNA-seq data to analyze Gankyrin (PSMD10) gene expression across various types of tissues and glioblastoma tumor samples. Processed RNA-Seq data for glioblastoma samples was downloaded from Cancer Genome Atlas TCGA as TPM from Xena portal [47]; normal TCGA samples and IDH mutant samples were excluded from the analysis. Next, for kept glioblastoma samples (N=147) we normalized library sizes and log-2 transformed expression profile.

Data for normal tissues was obtained from The Genotype-Tissue Expression (GTEx) on November 2024 (open access adult GTEX) [48]. The data was downloaded as TPM (Transcripts per million) and log-2 transformed to ensure comparability with TCGA data. Mann-Whitney U test was performed to identify differences in PSMD10 expression between glioblastoma samples and healthy tissues.

#### siRNA-mediated gene silencing in cell lines

Сells were seeded in a 12 well plate (100 000 cells per well) and in 24 hours were transfected for 24, 48, 72 and 96 hours with different concentrations of siRNA (1 nM, 5 nM, 10 nM) with Lipofectamine RNAiMax (Thermo Fisher Scientific, USA) transfection reagent following the protocol. Design, validation and selection of the most efficient siRNA for knockdown were described previously [12]. GBM23 and GBM24 were seeded in a 12-well plate at a density of 100,000 cells per well. After 24 hours, the cells were transfected with 10nM siRNA for 72 hours. Following this, a second transfection was performed for a further 72 hours.

#### Cytotoxicity assay of siRNA and its combination with drugs

*siRNA cytotoxicity.* Cells were seeded in a 96 well plate (3000 cells per well) and transfected for 24, 48, 72 and 96 hours with different concentrations of siRNA (1 nM, 5 nM, 10 nM), n=6. To determine siRNA cytotoxicity, 20 µl of Alamar Blue solution in culture medium was added to the cells and in 3 hours the 560/590 nm fluorescence intensity was measured using a VarioSkan Lux spectrophotometer (Thermo Fisher Scientific, USA). The percentage of living cells for each concentration of the substance was calculated by the formula (I - Ifon)/(Icontr - Ifon), where I is the average fluorescence intensity in the wells with cells; Icontr is the average fluorescence intensity in wells with control cells (untreated); Ifon is the average fluorescence intensity in wells with medium without cells.

*siRNA with cytotoxic drugs.* The cells (CT-2A, GL261, U87) were seeded in a 96-well plate (3,000 cells per well) and at the next day were transfected with siRNA to Gankyrin (5 and 10 nM) for 48 hours, siRNA to Luc and untreated cells were used as controls. Then the medium was replaced and cis or dox at the IC_50_ concentrations were added for each cell line for the subsequent incubation for 24 hours. In the case of TMZ the cells were incubated for 72 hours at different concentrations (250, 750, 1500, 2000 µM). The cytotoxicity of the cells was determined by the Alamar Blue test as described previously. Six technical repetitions were conducted for each drug concentration.

#### Western Blot

We extracted proteins with RIPA Lysis Buffer (Merck Millipore, USA) that contains a protease inhibitor cocktail (Roche, Switzerland). The bicinchoninic assay (BCA) was used to determine the total concentration of proteins in lysates. For that the following solutions were used: solution A: 8% solution of NaHCO3 and 1.6% solution of sodium tartrate adjusted to pH 11.25, solution B: 4% solution of BCA, solution C: 4% solution of CuSO4. The working solution was prepared in the ratio 25:25:1 of solution A:solution B:solution C. The standards of BSA (0-100 μg/ml) were used for the calibration plot. Proteins was heated at 95°C for 10 min and then in amount of 20 µg per lane were separated on 5%–12% SDS–polyacrylamide gels and transferred to PVDF membranes (Bio-Rad Laboratories Inc., USA) utilizing the Mini Trans-Blot® Cell (Bio-Rad Laboratories Inc., USA). Then the membranes were blocked in 3% milk, incubated with primary antibodies overnight at 4°C, and visualized using the appropriate HRP-linked secondary antibodies employing Clarity™ Western ECL Blotting Substrates (Bio-Rad Laboratories Inc., USA).

In our study, we employed several antibodies for immunological analysis. We utilized the mouse monoclonal Anti-β-Actin antibody (Sigma-Aldrich, #a1978) at a dilution of 1:5000. For Gankyrin detection, we used the anti-Gankyrin antibody (Abcam, #EPR14456(2)) diluted to 1:3000. For apoptosis detection anti-Caspase-1 (p20) monoclonal antibody (Casper-1) at a 1:3000 dilution was prepared (Adipogen, #AG-20B-0042-C100). Membranes were visualized using the appropriate HRP-linked secondary antibodies at a dilution of 1:10000 goat Anti-Mouse IgG (H+L)-HRP (Abcam, #ab205719) and goat Anti-Rabbit IgG (H+L)-HRP (BioRad, #1721019) - employing Clarity™ Western ECL Blotting Substrates (Bio-Rad Laboratories Inc., USA). Image quantification of the Western blot images was performed using ImageJ software (National Institute of Health, USA) in accordance with its standard procedure.

#### Reactive Oxygen Species (ROS) detection by 2’,7’-dichlorodihydrofluorescein diacetate (H2DCFDA)

Cells were seeded in 35mm confocal dishes (300 000 cells per well) and in 24 hours were transfected with siRNA to Luciferase and Gankyrin in a concentration of 10 nM for 24 hours. In order to detect reactive oxygen species (ROS) in cells after knockdown, unfixed cells were stained with H2DCFDA (Life Technologies, USA) for 15 min at 37°C with the replacement of culture medium with DMEM solution. The concentration of the dye in medium was 1 µM. After incubation cells were washed with DMEM two times for 3 minutes. Cell imaging was performed using a Nikon Eclipse Ti2 microscope. The final processing of the photos was performed by ImageJ software (National Institute of Health, USA).

#### Apoptosis/necrosis assay

Cells were seeded in 35mm confocal dishes (300 000 cells per well) and in 24 hours were transfected with siRNA to Luciferase and Gankyrin in a concentration of 10 nM for 24 hours. In order to detect apoptotic or necrotic signals in cells after knockdown, unfixed cells were stained with Apopxin Deep Red Indicator, CytoCalcein Violet 450 and Nuclear Green DCS1 Dye using the Abcam Apoptosis/Necrosis Assay Kit (Abcam, UK) following the manufacturer’s instructions or 45 min at 37°C with the replacement of culture medium with assay buffer. After incubation cells were washed with DMEM two times for 3 minutes. Cell imaging was performed using a Nikon Eclipse Ti2 microscope. The final processing of the photos was performed by ImageJ software.

#### Assessment of apoptosis induction using Caspase 3/7 assay

The induction of apoptosis was evaluated through the assessment of Caspase 3/7 activity with the Caspase-Glo® 3/7 assay kit (Promega, USA). The method employed a luminescent detection based on the luciferin-luciferase system.

The U87 cell line was cultured in 96-well plates at a density of 10,000 cells per well. Following a 24-hour period, the cells were transfected with 10 nM siRNA for a further 48 hours, with 1% DMSO employed as a positive control. Subsequently, the Caspase-Glo® 3/7 reagent was prepared in accordance with the manufacturer’s instructions. Then, 25 μL of the reagent was added to each well, and the plates were incubated for 2 hours at room temperature. Luminescence was measured using a microplate reader Varioskan Lux afterward.

#### Cell Cycle Analysis with Flow Cytometry

GL261 and U87 cells were seeded in 6-well plates in triplicate (approximately 300 000 cells per well) and transfected with siRNA at 2 concentrations (5nM, 10 nM). In 48 hours after transfection cells were collected, washed twice with PBS and fixed overnight at 4 ◦C with 2 mL of 70% ethanol. After fixation, cells were washed with PBS and then re-suspended in 0.5 mL of PBS with 5 µg/mL RNase A and 30 µg/mL PI (propidium iodide). Cell cycle measurement was performed by Flow Cytometer BD FACSCanto. The flow cytometry results were analyzed using FlowJo™ v10.8 Software (BD Life Sciences, Franklin Lakes, NJ, USA).

#### Stereotaxic implantation of GL261 cells

All experiments were approved and performed in accordance with institutional guidelines approved by the Animal Ethics Committee of Pirogov Russian National Research Medical University.

Mice (C57Bl6) were deeply anesthetized with 3% isoflurane (Baxter, USA) in an induction chamber using a low-flow anesthesia system (SomnoSuite, Kent Scientific). Then, mice were placed onto the heating pad of the stereotaxic apparatus (digital stereotaxic instrument, Stoelting Co). Anesthesia was maintained by delivery of 1.5% isoflurane via inhalation mask. Hair on the head was shaved and disinfected with ethanol. Longitudinal incision was made on the skull with a scalpel, and the surface of the skull was treated with hydrogen peroxide using cotton buds and dried by H2O2 to facilitate identification of the bregma. The skull surface was aligned in anteroposterior and mediolateral directions. GL261 cells in 50.000 cells in 5 μl were injected into striatum using a Hamilton microsyringe with coordinates (AP -1.5 mm; ML -1.0 mm; DV -3 mm). A syringe with implantable cells was injected to the -3.5 mm and waited 3 minutes for the tissues to diverge. After that the syringe was raised by 0.5 mm and the cell solution was injected at a rate of 500 nl/min. To stabilize the pressure in the tissues it was waited for 3 minutes. The hole was closed with a filling solution, irradiated with ultraviolet light for 20 seconds. Finally, the skin wound was sutured with surgical silk suture.

#### Magnetic resonance imaging (MRI)

The tumor growth of glioma cells (GL261 or GBM23) cells in mice was studied by scanning in T2 Turbo Spin Echo mode (TE/TR=46/3720 ms, slice thickness 0.5 mm, resolution 384/288 for the transversal plane) on a ClinScan 7T magnetic resonance tomography (Bruker Biospin, USA) at 7, 14, and 21 days (for GL261) and 1 and 3 month (for GBM 23) after cell implantation.

#### In vivo accumulation of LNP-siRNA

For biodistribution experiments we intratumorally and intravenously injected cyanine 5 formulated into LNP (5 μl, 130 ng/μl) to the mice with implanted GL261 cells for 3 weeks and analyzed the whole body fluorescence using IVIS Spectrum CT (Perkin Elmer) in 3, 6, 12 and 24 hours after injection at 640/680 excitation/emission filter. In 24 hours after injection mice were anesthetized (Zoletil 20 mg/kg, xylazine 0.2 mg/kg) and perfused with 20 ml of PBS and then with 30 ml of 4% formalin. The brain, lungs, heart, liver, kidneys and spleen were isolated and analyzed for fluorescence using IVIS Spectrum CT at 640/680 excitation/emission filter. After that, the brains were fixed in 4% paraformaldehyde for 24 hours and cut using vibratome (Campden Instruments, England). The imaging of 50 μm slices was performed using a Nikon Eclipse Ti2 microscope. The fluorescence of GFP and cyanine 5 formulated into LNP was analyzed at 488/509 and 640/680 excitation/emission filters.

#### In vivo Gankyrin knockdown

To perform in vivo knockdown of Gankyrin, we injected GL261 cells (50 000 cells/ 5µl) into striatum and then in 3 weeks after implantation we intratumorally administered NaCl or siRNA (siLuc was used as a control) formulated into LNP (1 µg/5 µl). In 72 hours after injection mice were anesthetized (Zoletil 20 mg/kg, xylazine 0.2 mg/kg) and small pieces of tumor and healthy brain were isolated for RNA extraction, cDNA synthesis and evaluation of Gankyrin gene expression by RT-PCR.

#### Cultivation of patient-derived glioma cells, validation of GBM model and siGankyrin treatment

Clinical glioblastoma specimens were collected in the Neurosurgery department оf Federal Center of Brain research and Neurotechnology of the Federal Medical-Biological Agency of Russia with informed patients consent and processed to the research laboratories after de-identification of the samples. All diagnoses were confirmed by morphological studies. The study was approved by the ethics committees of the Federal Center of Brain research and Neurotechnology of the Federal Medical-Biological Agency of Russia. Glioblastoma specimens were collected during surgery and mechanically dissociated into pieces with 1-3 mm diameter. The samples were then treated with collagenase II for 20 minutes at +37 °C to obtain single cells. Established cell lines were cultivated in serum-free DMEM/F12 media, supplemented with Neuromax (Paneco, Russia) 20 ng/ml basic fibroblast growth factor (bFGF; Sci-Store), and 20 ng/ml epidermal growth factor (EGF; Sci-Store).

The primary endpoints were to assess tumor growth inhibition after siGankyrin administration and corresponding survival rates of the treated mice. GBM xenograft models were initiated in 8-week old NOD/SCID mice housed under pathogen-free conditions. A maximum of five mice were housed per cage in a controlled environment with a 12-hour light/dark cycle, temperature maintained at 22 °C, and free access to food and water. A total of 8 NOD/SCID mice were enrolled in the study, with a balanced randomization resulting in two groups: one receiving intratumoral siGankyrin (n=4), one receiving a control treatment (PBS, n=4). Mice were excluded if they exhibited any signs of significant distress or morbidity prior to the initiation of the experiment, or if they experienced any complications from the surgical procedures, such as infection or prolonged recovery times. For intracranial cell injections 500,000 GBM23 cells in 2 μl of PBS were injected 3 mm below the brain surface, 1.5 mm rostral of the bregma and 1.0 mm right of the midline, using a 30G Hamilton needle and a 2 μl syringe in a stereotaxic frame. Mice were anesthetized with 2% isoflurane in oxygen at a flow rate of 2 ml/min. The burr hole through which the cells were injected was sealed with bone wax and the midline scalp incision was closed with silk suture. Mice were euthanized when they exhibited signs of significant morbidity (hunching, weight loss, rough coat, ataxia, head tilt, paralysis). After one month of the implantation of GBM23, siGankyrin was administered intratumorally at 72-hour intervals, with four administrations of 1 μg each. Outcome measures included overall survival time, which was recorded from the date of initial tumor implantation until the date of euthanasia due to significant morbidity. Survival analyses were conducted using log-rank tests for comparisons between groups.

All studies were conducted according to protocols approved by the Animal Ethics Committee of Pirogov Russian National Research Medical University.

#### Statistical Analysis

All diagrams presented here are based on at least three independent experiments. Statistical processing of obtained data was performed using the GraphPad Prism software (version 8.0.1) (GraphPad Holdings, LLC, San Diego, CA, USA) with multiple t-tests and one-way analysis of variance (ANOVA). The data were considered statistically significant at p-value < 0.05. Comparison of survival curves was prepared with Comparison of Survival Curves log-rank (Mantel-Cox) test.

---

**Figure S1.**
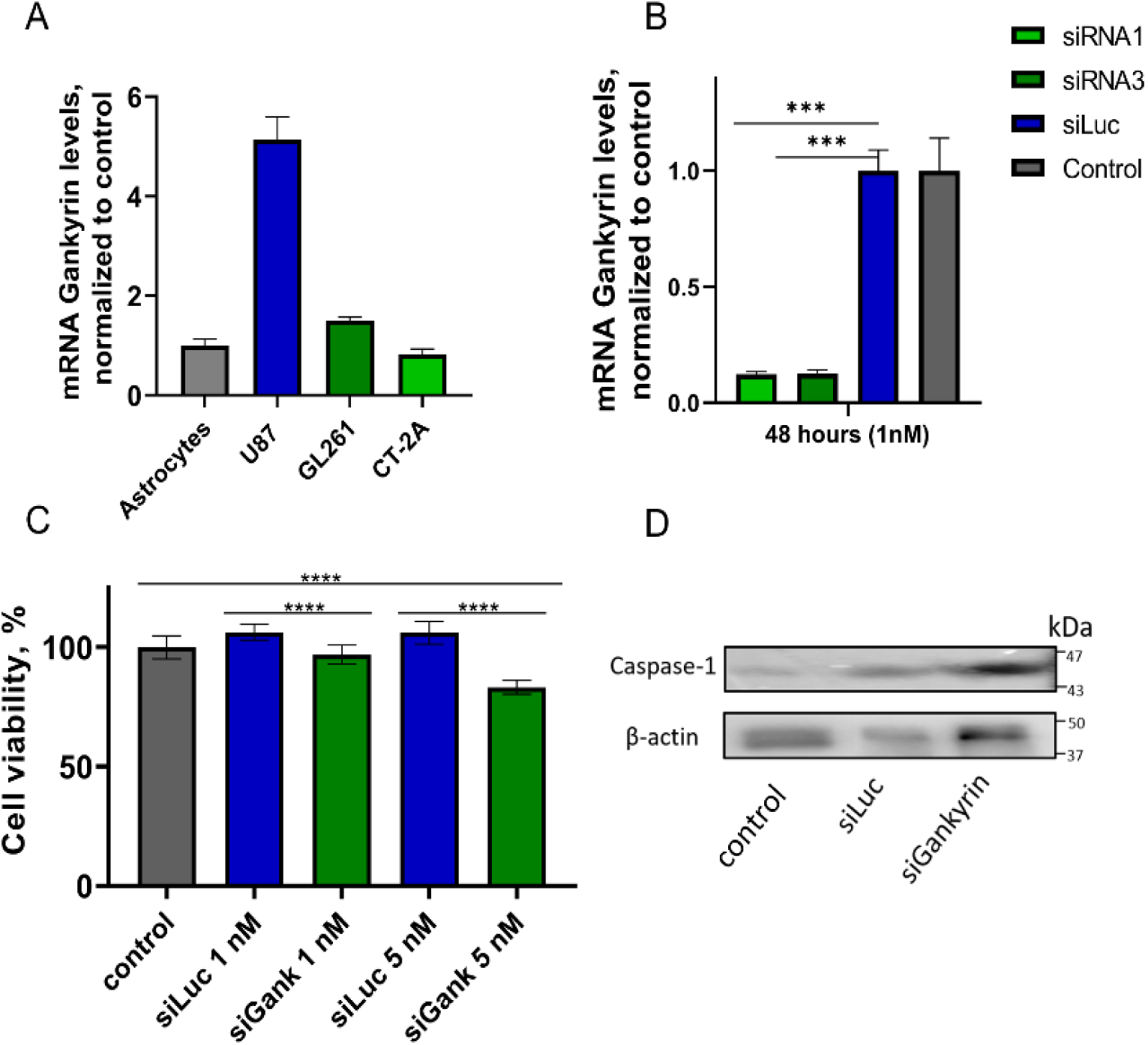
Gankyrin inhibition and its effects. (A) Gankyrin expression in different cell lines. Astrocytes were used a a control. (B) Gankyrin expression in U87 cells after 48 hours of transfection with 1 nM siRNA. (C) Cell viability after transfection with siRNA. GL261 viability in 72 hours after transfection with siRNA to Gankyrin in 2 concentrations (1 nM, 5 nM) compared to the control. (D) Western blot analysis of caspase-1 (p20) expression in GL261 48 hours after knockdown with siGankyrin (10nM)

**Figure S2.**
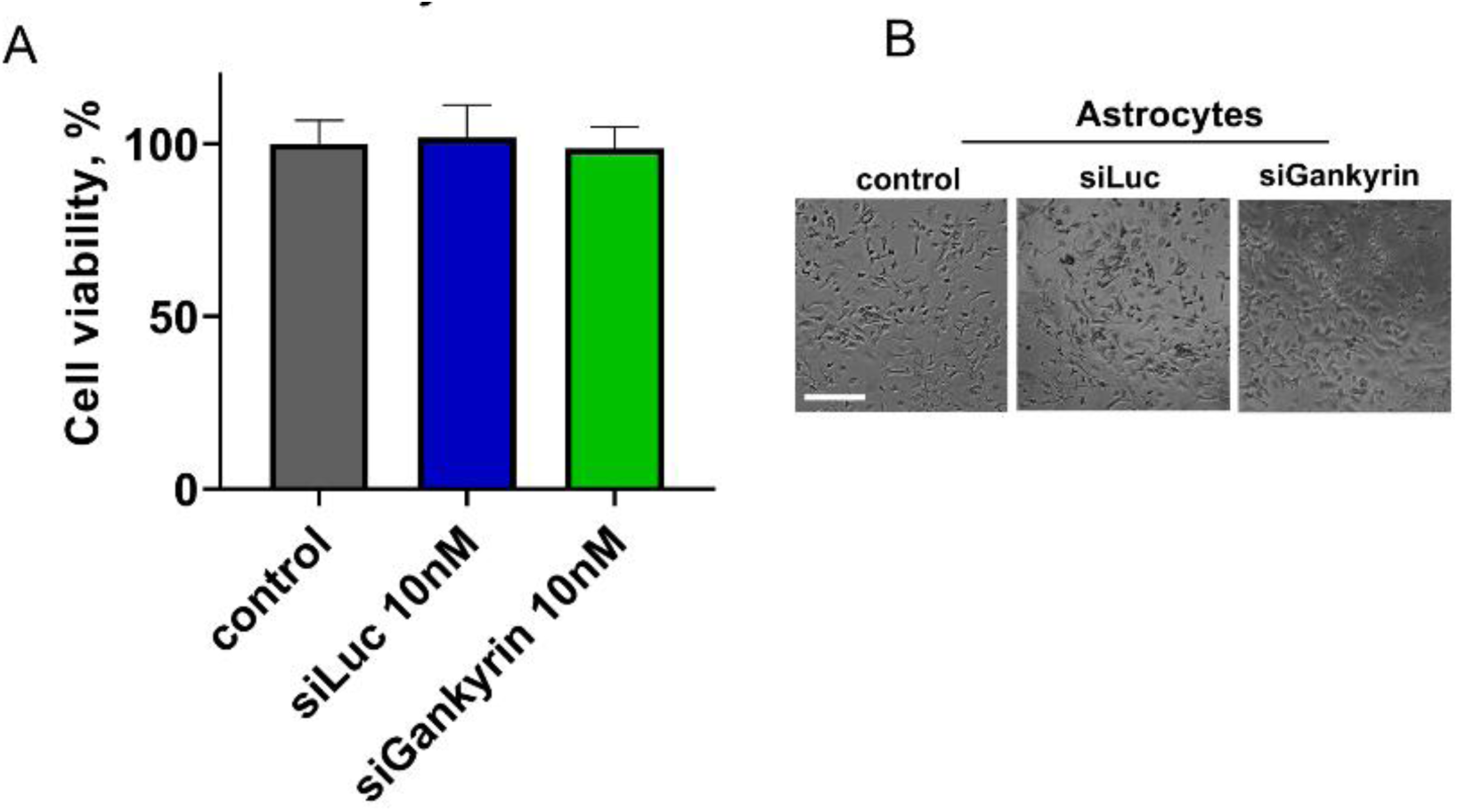
Cell viability after transfection with siRNA. (A, B) Murine astrocytes viability in 48 hours respectively after transfection siRNA. Scale bar 50 µm. Five biological replicates per sample were used; results represented mean ± SD. (** p < 0.01,*** p < 0.001, **** p < 0.001, One-way ANOVA)

**Figure S3.**
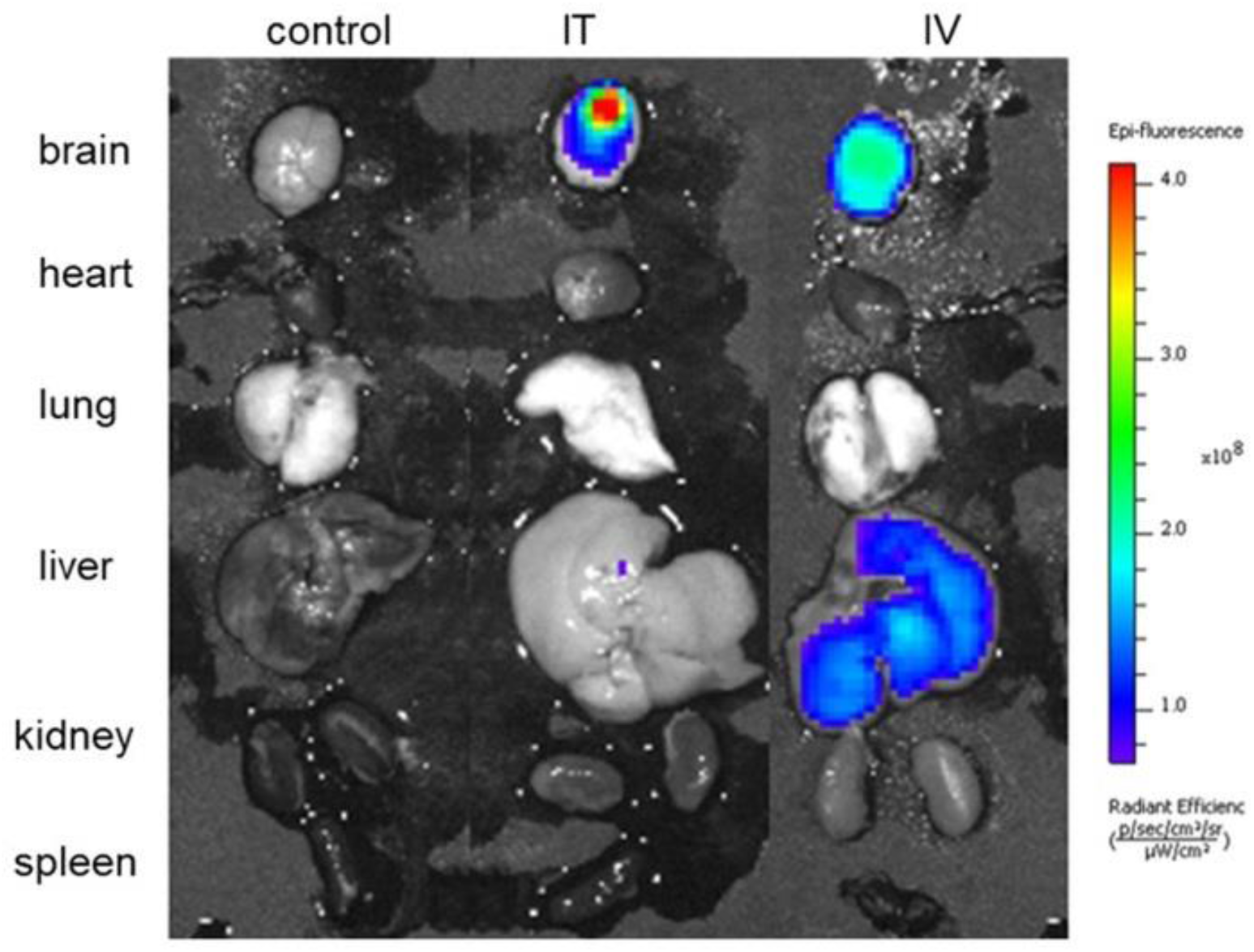
Biodistribution of siRNA-LNP after intratumoral (IT, early-stage glioma) and intravenous (IV, late-stage glioma) injection to mice with orthotopic GL261 glioma model in different organs in 24 hours.

**Table S1.** Half inhibitory concentration of 3 drugs (doxorubicin, cisplatin, temozolomide) in different cell lines (µM)

| Cell line | IC<sub>50</sub> doxorubicin, $\mu\text{M}$<br>(48h) | IC<sub>50</sub> cisplatin, $\mu\text{M}$<br>(48h) | IC<sub>50</sub> temozolomide, $\mu\text{M}$<br>(72h) |
| --- | --- | --- | --- |
| U87 | 0,18±0,12 | 12,3±1,7 | 900±140 |
| GL261 | 0,47±0,04 | 9,0±1,2 | 21±8 |
| CT-2A | 0,17±0,07 | 25,8±1,3 | 1200±90 |
| GBM23 |  |  | 1080±110 |
| GBM24 |  |  | 770±115 |

